# Auditory spatial analysis in reverberant audio-visual multi-talker environments with congruent and incongruent visual room information

**DOI:** 10.1101/2022.04.30.490125

**Authors:** Axel Ahrens, Kasper Duemose Lund

## Abstract

In multi-talker situation, listeners have the challenge to identify a target speech source out of a mixture of interfering background noises. In the current study it was investigate how listeners analyze audio-visual scenes with varying complexity in terms of number of talkers and reverberation. Furthermore, the visual information of the room was either coherent with the acoustic room or incoherent. The listeners’ task was to locate an ongoing speech source in a mixture of other speech sources. The 3D audio-visual scenarios were presented using a loudspeaker array and virtual reality glasses. It was shown that room reverberation as well as the number of talkers in a scene influence the ability to analyze an auditory scene in terms of accuracy and response time. Incongruent visual information of the room did not affect this ability. When few talkers were presented simultaneously, listeners were able to quickly and accurately detect a target talker even in adverse room acoustical conditions. Reverberation started to affect the response time when four or more talkers were presented. The number of talkers became a significant factor for five or more simultaneous talkers.

## I. Introduction

The human auditory system has the ability to focus on a speech stream in the presence of interfering speech stimuli. Such a multi-talker scenario has been termed the cocktail-party situation (Bronkhorst, 2000; Cherry, 1953). Many factors are known to reduce the ability to understand speech in such a cocktail-party situation, e.g., the level of the target speech relative to the interferers, the number of talkers, or the type of listening room. These effects are commonly measured by asking the listeners to repeat a word or a sentence or to write down the perceived stimulus. However, in our daily life the task in a cocktail-party situation is usually different, where it is necessary to follow a conversation and to identify a certain topic or continuous speech stream out of an interfering speech mixture. In the current study we investigated the ability of listeners to analyze an acoustic scene with varying complexity in terms of number of interfering talkers, room reverberation and coherency of visual room information.

The number of interfering talkers has been shown to influence the intelligibility of a target talker. (S. A. Simpson & Cooke, 2005) showed that the intelligibility decreases when increasing the number of interfering speech sources for up to eight interfering talkers, as the ability to listen into speech gaps is reduced and at the same time the interfering speech remains intelligible and can be confused with the target speech. When further increasing the number of interfering talkers, the intelligibility was shown to improve as the interferers become more noise-like and therefore do not contain understandable speech.

Reflections and reverberation are present in nearly all communication scenarios. Room reverberation has been shown to negatively affect speech perception in a number of studies (Best et al., 2015; Bronkhorst & Plomp, 1990; Moncur & Dirks, 1967; Nabelek & Mason, 1981; Nábělek & Pickett, 1974). Particularly, the diffuse reverberation, i.e., the late reverberant tail, has been shown to reduce speech intelligibility, while early reflections do not seem to harm, or might even improve speech perception (Arweiler et al., 2013; Arweiler & Buchholz, 2011; Warzybok et al., 2013).

Previous studies have investigated the ability of listeners to identify and locate speech in the presence of other speech sources. (Kopčo et al., 2010) measured the localization accuracy of a digit spoken by a female talker in the presence of words spoken by male interfering talkers. The target and the interferers were all presented in the frontal area of the listener. They found that the presence of the interferers reduced the localization accuracy. (Buchholz & Best, 2020) measured localization accuracy with a similar target digit as in (Kopčo et al., 2010) but with a more realistic background noise scene. The interfering signals were seven paired conversations (both male and female) at various locations in a simulated cafeteria. Results showed that the localization accuracy was only affected by the noise when the target source was distant but not when it was nearby. This finding suggests an interaction with reverberation, as farther sources have more reverberant energy relative to the direct sound compared to nearby sources.

While these studies focused on the ability to locate a speech signal in a speech background, (Hawley et al., 1999) investigated both the localization accuracy of speech as well as the intelligibility. They showed that the inability to correctly locate a source did not limit the ability to correctly understand it. However, the number of interfering sources was limited to three. (Weller et al., 2016) presented a novel method to evaluate the ability to analyze a complex acoustic scene. They asked their listeners to judge the location of all talkers presented in a virtual cocktail-party situation by indicating the gender of the talkers. When varying the number of simultaneously presented talkers, they found that normal-hearing listeners were able to correctly locate and count the number of talkers for up to four sources. When six talkers were presented, the accuracy decreased.

Most of the beforementioned studies focused on the ability to localize speech but less to comprehend the speech. However, in a real-world cocktail party, listeners need to perform both tasks to successfully communicate. In the current study, we asked listeners to locate a talker speaking about a certain topic, while presenting a varying number of other simultaneous talkers. Thus, the primary task was to understand the speech and the secondary task to locate the talker. The experiment was conducted in an audio-visual virtual environment using a loudspeaker array and virtual reality glasses. The listeners’ task was to indicate a semi-transparent avatar at the location of an acoustic source talking about a topic indicated by an icon. The sources were located at one of fifteen possible locations with 15° horizontal separation. The number of simultaneous speech sources was varied between two and eight. Three virtual rooms were simulated visually and acoustically. Furthermore, a condition with incongruent audio-visual cues was presented by visually showing the anechoic room and acoustically presenting the reverberant room or vice versa.

## II. Methods

### A. Participants

Thirteen Danish native speaking normal-hearing listeners aged 20-26 years participated in the experiment (7 female and 6 male). Participants were paid on an hourly basis and gave consent to an ethics agreement approved by the Science-Ethics Committee for the Capital Region of Denmark (reference H-16036391).

### B. Material

The speech material for target and interferers was taken from a database of anechoically recorded monologues in Danish (see (Lund et al., 2019) for details^1^). Each monologue was designed with characteristic features in mind, ensuring significant difference of the content. The database consists of ten monologues each spoken by ten native Danish speakers.

### C. Audio-visual rooms

Three different acoustic and visual rooms were used in this study, a high-reverberant room, a mid-reverberant room and an anechoic room. The dimensions of all three rooms remained constant as shown in Figure 1, both acoustically and visually. However, the surface materials differed.

**Figure 1:**
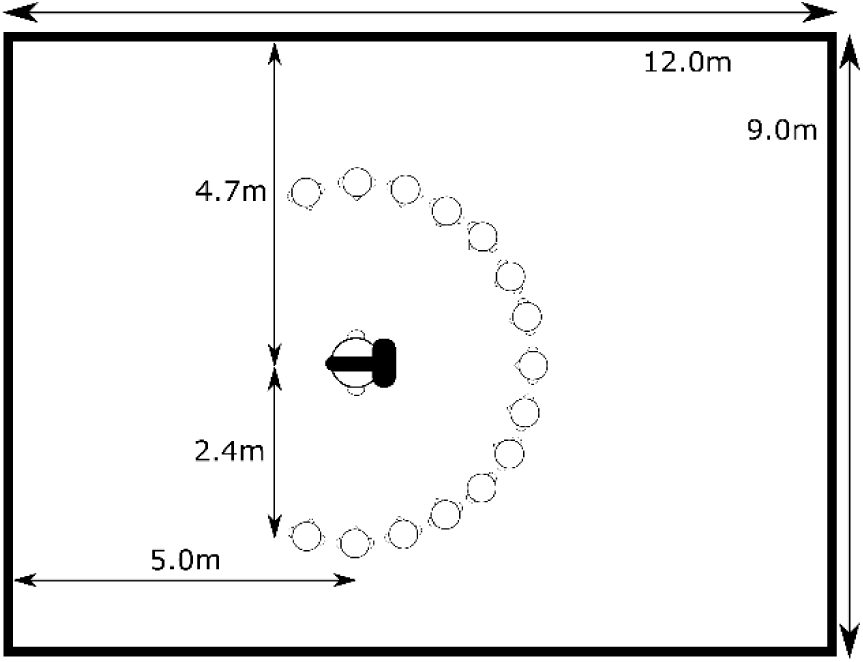
Top view of the virtual audio-visual room. The listener is wearing VR glasses with a visual simulation of the room including 15 potential talker positions at 2.4m distance in the frontal hemisphere visualized by the head icons. The height of the room is 2.8m.

Figure 2 shows the visual appearances of the three rooms. Figure 2A shows the anechoic room with foam wedges as commonly seen in anechoic chambers. For the acoustic reproduction of this room only the direct sound was reproduced from single loudspeakers. In Figure 2B the mid-reverberant room can be seen. The visual as well as the acoustical properties were similar to a large living room. The highly reverberant room is shown in Figure 2C. It was modelled with bare concrete surfaces to simulate a highly reverberant, yet realistic environment.

**Figure 2:**
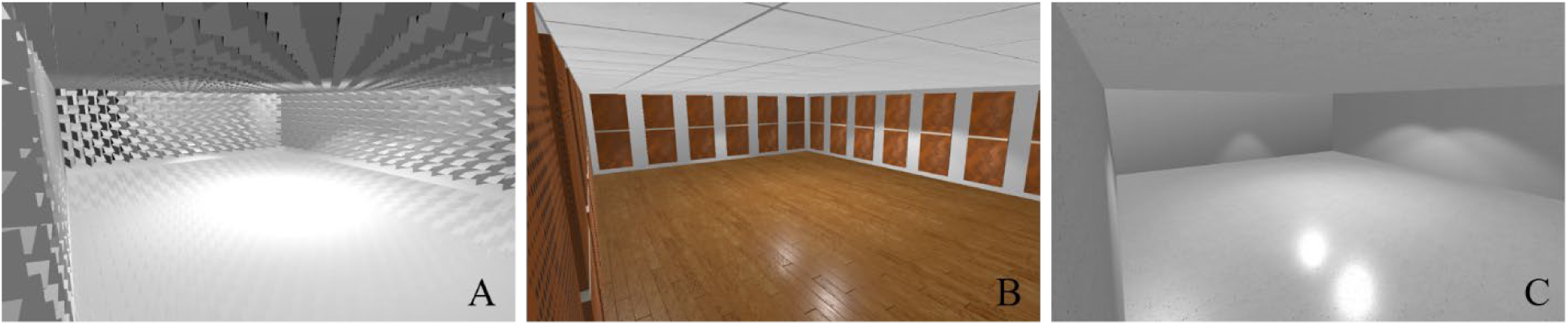
Visual appearance of the three virtual rooms. A: anechoic, B: mid-reverberant, C: high-reverberant. The dimensions in the rooms are identical, while the surface materials differ.

The rooms were simulated using the room acoustic simulation software Odeon (Odeon A/S, Kgs. Lyngby, Denmark) with the materials and surface absorption coefficients as shown in Table 1. For the anechoic room, only the direct sound was considered. In Figure 3 the reverberation time, clarity and direct-to-reverberant ratio of the three rooms are shown. The reverberation time as well as the clarity were calculated using the ITA-toolbox (Berzborn et al., 2017), the direct-to-reverberant ratio was calculated as the ratio between the direct sound and the reflections. Mind that for the anechoic condition the clarity and direct-to-reverberant ratio are infinite as no reflections are present which is indicated with arrows.

**Table 1:** Absorption coefficients (α) of the surfaces in the mid-reverberant and high-reverberant room.

| $\alpha$ (mid-rev/high-rev) | 63 Hz | 125 Hz | 250 Hz | 500 Hz | 1 kHz | 2 kHz | 4 kHz | 8 kHz |
| --- | --- | --- | --- | --- | --- | --- | --- | --- |
| Side walls<br>Wooden panels/Brick | 0.2/0.06 | 0.2/0.06 | 0.2/0.06 | 0.3/0.07 | 0.4/0.07 | 0.4/0.07 | 0.5/0.08 | 0.5/0.09 |
| Floor<br>Parquet/Concrete | 0.2/0.05 | 0.2/0.05 | 0.15/0.05 | 0.1/0.05 | 0.1/0.07 | 0.05/0.07 | 0.1/0.07 | 0.1/0.07 |
| Ceiling<br>Gypsumboard/Concrete | 0.3/0.05 | 0.3/0.05 | 0.35/0.05 | 0.4/0.05 | 0.4/0.07 | 0.4/0.07 | 0.5/0.07 | 0.55/0.07 |

**Figure 3:**
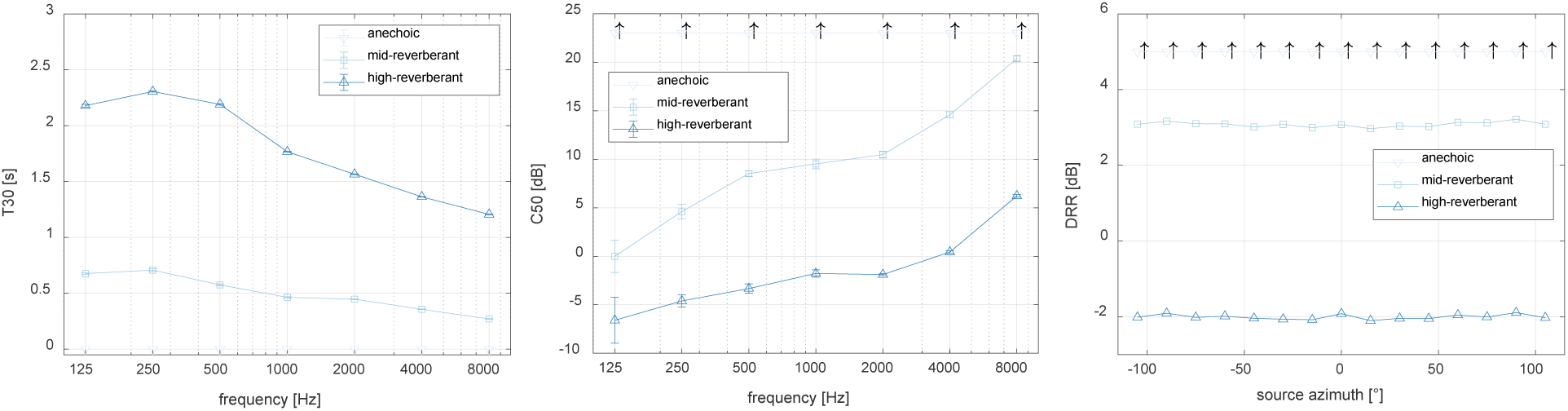
Reverberation time (T30), Clarity (C50) and the direct-to-reverberant ratio (DRR) for the three rooms. The T30 and the C50 are shown with respect to octave frequency bands. The DRR is shown with respect to the source azimuth angle. The arrows indicate that the measure is infinite.

### D. Task

The listeners’ task was to identify the location of a talker amongst concurrent talker(s) in a virtual audio-visual room according to the story in the monologue. Accuracy and completion time of the task was emphasized by advising the listeners to “find the correct story as fast as possible”. The number of concurrent talkers varied between two and eight, thus the number of interfering talkers varied between one and seven. An icon visualizing the target story content was displayed on the backwall in the visual virtual room. The 15 possible talker positions were always represented by semi-transparent humanoid shapes independent of the actual number of concurrent talkers. Figure 1 visualizes the possible talker locations between −105° to 105° separated by 15° in the frontal hemisphere at a distance of 2.4 m. The task was performed by pointing at the position where the target talker was perceived. The participants were using a virtual reality controller that included the visual appearance of a laser pointer in the virtual room.

For each scene a unique talker, story and position was randomly chosen as the target. Between one and seven masking talkers were included in a similar way. No talker, story or position could occur twice at the same time. For each trial, the acoustic talkers were presented for 120 seconds. The stories were started at a random point in time and were repeated from the beginning after finishing. Thus, no bias towards the beginning of each story was introduced. The listener could indicate the perceived target talker position at any time, even after the audio had stopped. Each individual talker was presented at a sound pressure level of 55 dB SPL.

Three congruent audio-visual rooms were used as described above, an anechoic, a mid-reverberant and a high-reverberant room. In addition to the conditions with congruent audio and visual room information, two conditions with incongruent audio-visual cues were considered. These were anechoic acoustics with the appearance of a highly reverberant room and high-reverberant acoustics with the visuals of the anechoic room. Thus, five room conditions were tested. Each of the conditions was repeated three times resulting in 105 trials, five audio-visual conditions and between two and eight concurrent talkers.

Prior to the experiment, the listeners performed a familiarization phase, where they were familiarized with the speech material and the story content but not with the task itself. The anechoic version of the ten stories were played back via headphones in a randomized order. Each talker was randomly assigned to one of the stories. Thus, listeners heard each story and each talker once. For the training, listeners were instructed to focus on unique content features or passages of the stories. After completed training listeners were seated in the loudspeaker environment and introduced to the listening task and the interaction method using the VR controller.

### E. Virtual audio-visual setup

The virtual visual scenes were rendered on the head-mounted display (HMD) of an HTC Vive Pro Eye (HTC Vive system, HTC Corporation, New Taipei City, Taiwan). This system allowed to track the listeners motion and record eye gaze and pupil dilation from inside the HMD with a sampling frequency of up to 120 Hz and an accuracy between 0.5° and 1.1°. The visual virtual scenes were modeled and displayed using Unity (Unity Technologies, San Francisco, California, USA).

The acoustic scenes were reproduced on 64-channel spherical loudspeaker array housed in an anechoic chamber (see (Ahrens, Marschall, et al., 2019) for details). The loudspeaker signals were generated using the room acoustic simulation using the LoRA-toolbox (Favrot & Buchholz, 2010). For the loudspeaker playback the nearest loudspeaker mapping was applied, where the direct sound as well as the early reflections are mapped to the nearest loudspeaker. The late reverberant tail is reproduced using 1^st^ order ambisonics to achieve a diffuse acoustic field (Favrot & Buchholz, 2010).

### F. Outcome measures and statistical analyses

To evaluate the listeners’ ability to successfully analyze a cocktail-party scenario, two outcome measures were evaluated. First, the ability to correctly identify and locate the target talker. This allows for a binary right/wrong analysis as well as a localization error in degrees. Second, the response time of the listener from audio onset to decision.

The outcome measures were analyzed using an analysis of variance of mixed linear models. The computational analyses were done using the statistical computing software R(R Core Team, 2020) and the lmerTest (Kuznetsova et al., 2017) package. Within factor analyses were conducted using marginal means implemented in the emmeans package (Lenth, 2020) with Tukey correction for multiple comparisons.

## III. Results

### A. Coherent audio-visual room information

Figure 4 shows the percentage of correctly located stories. Each bar contains 42 datapoints across the 14 participants and three repetitions. When few talkers are in a scene, the participants were able to accurately locate the correct story in all reverberation conditions. In scenes with more than five talkers, the accuracy in the high-reverberant condition (dark blue) decreases. In the mid-reverberant condition such a decrease can only be observed when eight talkers are in a scene. In the anechoic condition, the participants were able to accurately locate the target story for all numbers of talkers.

**Figure 4:**
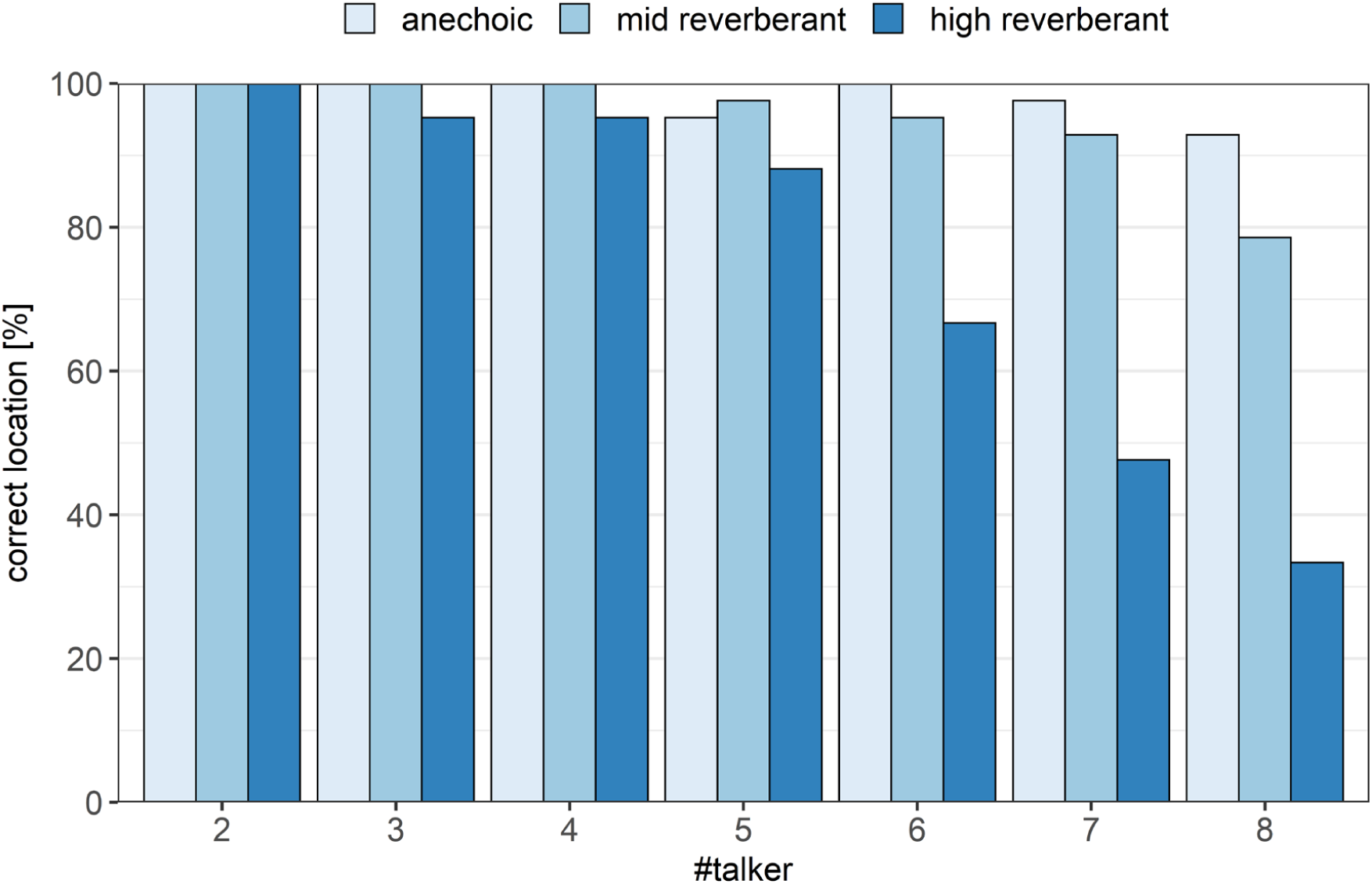
The percentage of correct response locations. Each bar contains 42 datapoints across subjects and repetitions. The three colors indicate the room conditions.

Figure 5 shows the response time of the correct responses when two to eight talkers were presented simultaneously. The response time is displayed for the audio-visually coherent room conditions with varying reverberation times indicated with the different colors. With an increasing number of simultaneous talkers, the time needed to identify the target talker increased [F(6,755.2)=73.1, p<0.0001]. The response time was also found to be dependent on the reverberation time [F(2,755.6)=83.1, p<0.0001]. Furthermore, the interaction term between the number of talkers and the reverberation time was found significant [F(12,754.8)=5.4, p<0.0001]. Specifically, the high-reverberant condition was found to lead to a higher response time when four or more talkers were presented [p<0.05] but not with less than four talkers [p>0.5]. The differences between the high-reverberant condition and the anechoic/mid-reverberant condition increases with larger numbers of talkers. No significant differences between the anechoic and the mid reverberant condition was found [p>0.1] across all number of talkers.

**Figure 5:**
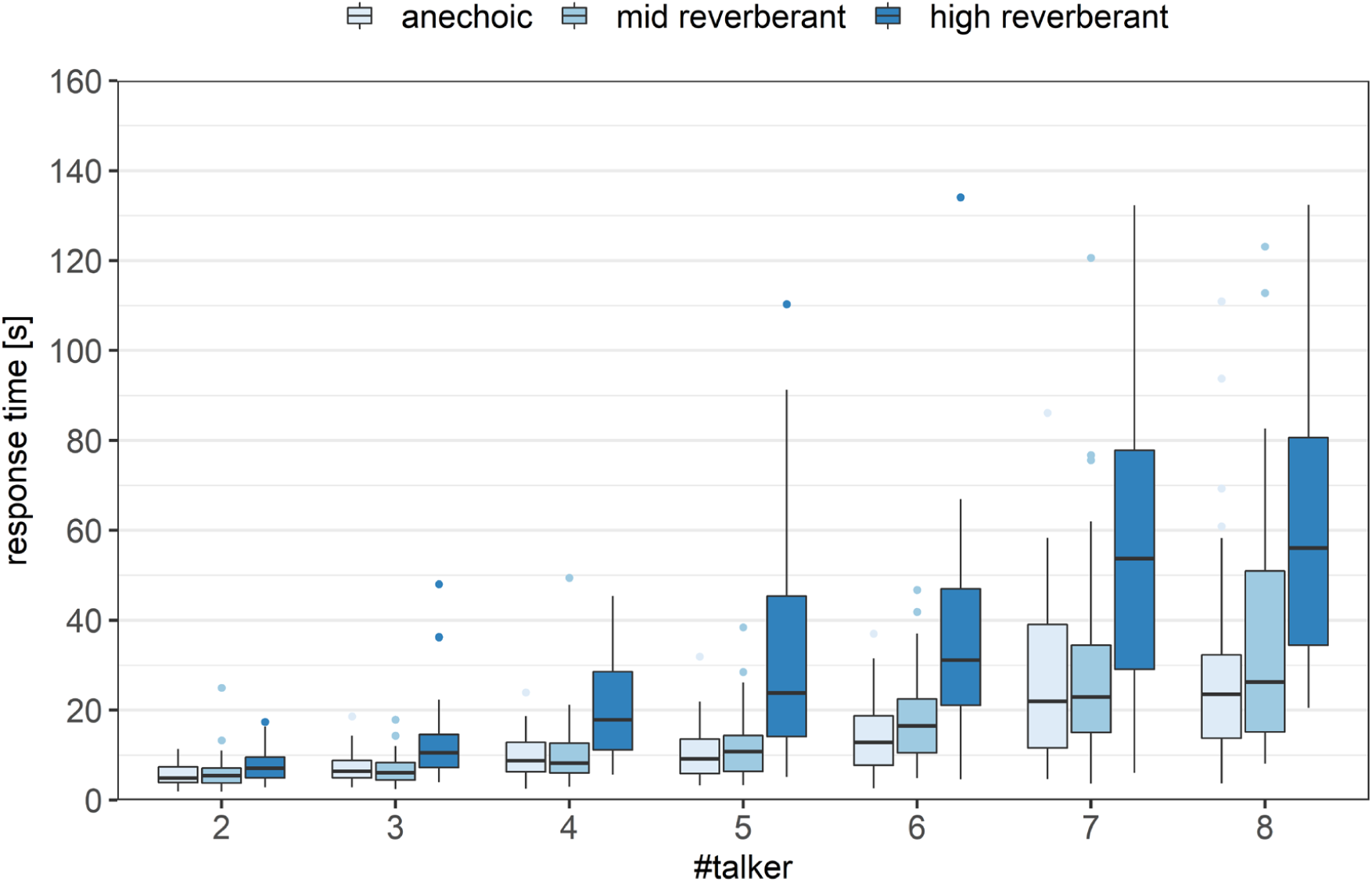
Response time with respect to the number of talkers in a scene of all correct responses. The colors indicate the room reverberation conditions. The boxes cover the range between the 25th and the 75th percentile. The horizontal line in the boxes indicates the median. The whiskers extend to 1.5 times the inter-quartile range. Outliers are indicated as dots.

In Figure 6 the localization error is shown. In the high-reverberation condition an increasing mean localization error was found for six and more talkers, with the eight-talker setting resulting in a median error of 30°, i.e., two potential positions error from the target location. In the anechoic and mid-reverberant conditions only few errors were found, indicated as outliers in Figure 6.

**Figure 6:**
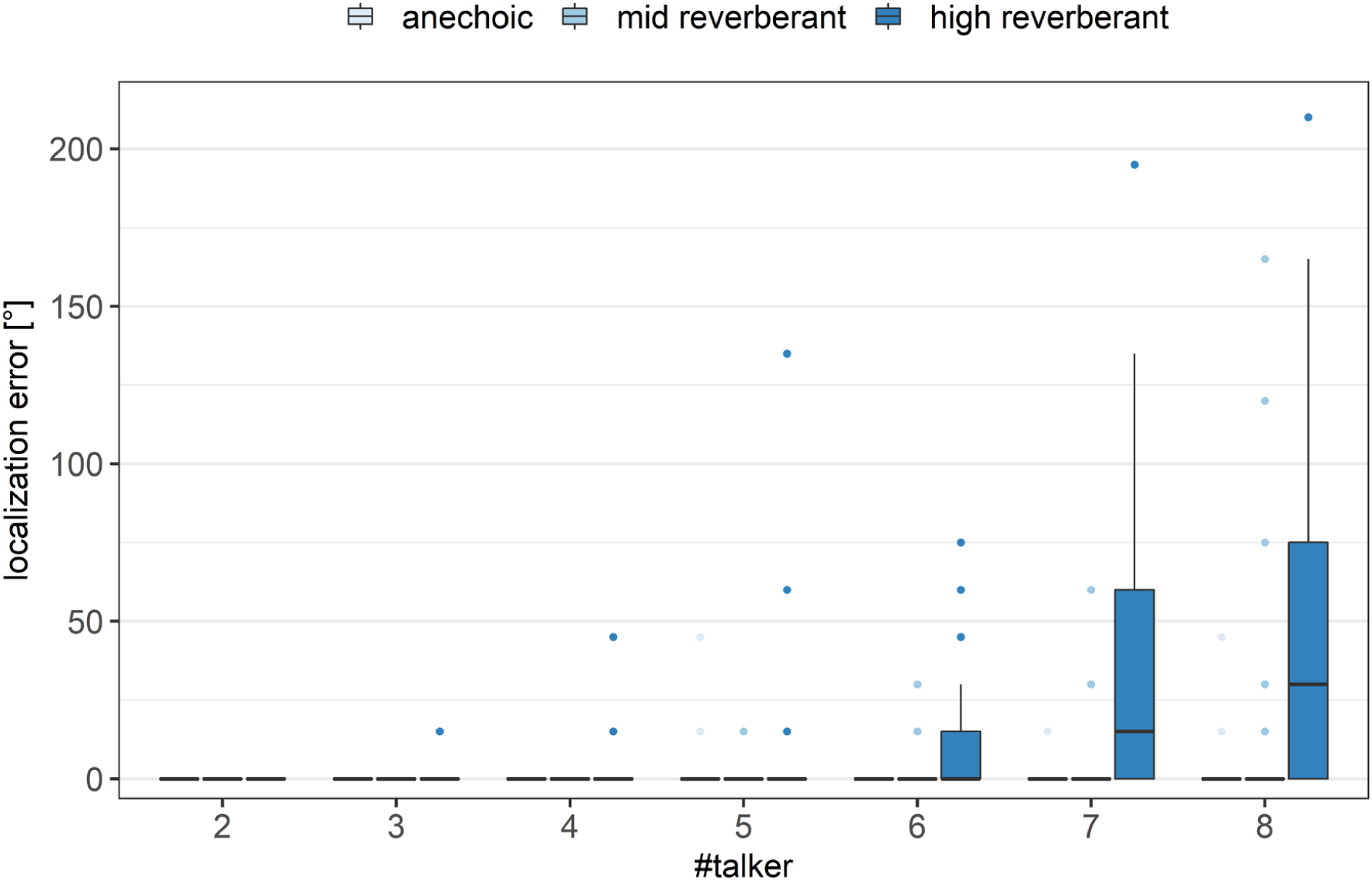
Localization error with respect to the number of talkers. The three colors indicate the room conditions. The boxes cover the range between the 25th and the 75th percentile. The horizontal line in the boxes indicates the median. The whiskers extend to 1.5 times the inter-quartile range. Outliers are indicated as dots.

### B. Incoherent audio-visual room information

Figure 7 shows the percentage of correctly identified stories, comparing the coherent and the incoherent audio-visual conditions with and without reverberation. The light blue/grey bars indicate the acoustically anechoic conditions and the dark bars the acoustically reverberant conditions. No differences arise from the audio-visual incongruency.

**Figure 7:**
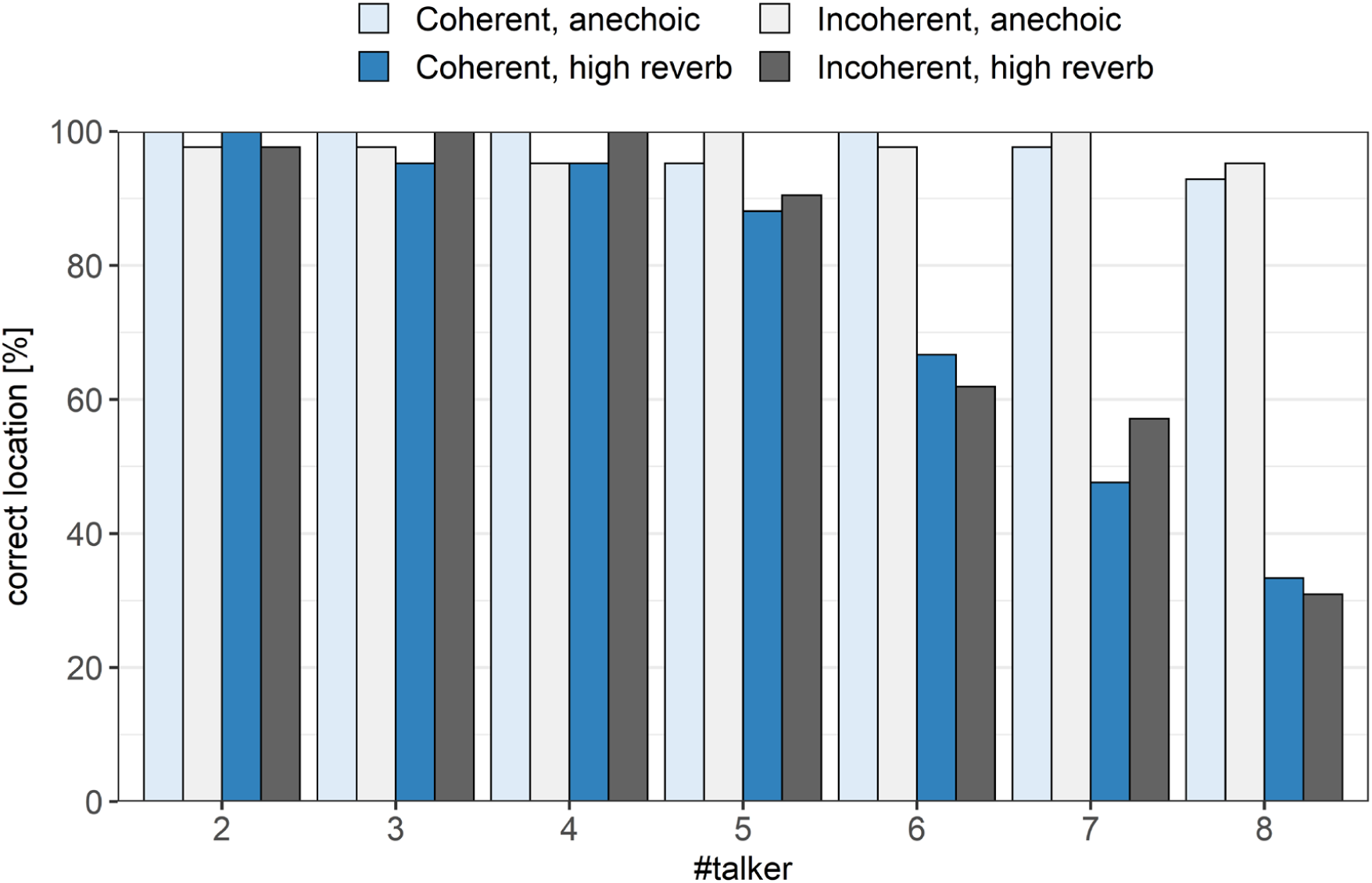
The percentage of correct response locations comparing the coherent and incoherent audio-visual conditions. Each bar contains 42 datapoints across subjects and repetitions. The three colors indicate the room conditions.

Figure 8 shows the response times for the incongruent audio-visual conditions (grey boxes), i.e., the conditions with anechoic acoustic stimuli and the visuals of the reverberant room (light grey) and with high acoustic reverberation and the visuals of the anechoic room (dark grey). Additionally, the response times from the coherent anechoic and reverberant conditions are shown (blue boxes, as in Figure 5). No significant difference was found between the congruent and the incongruent condition [p>0.12].

**Figure 8:**
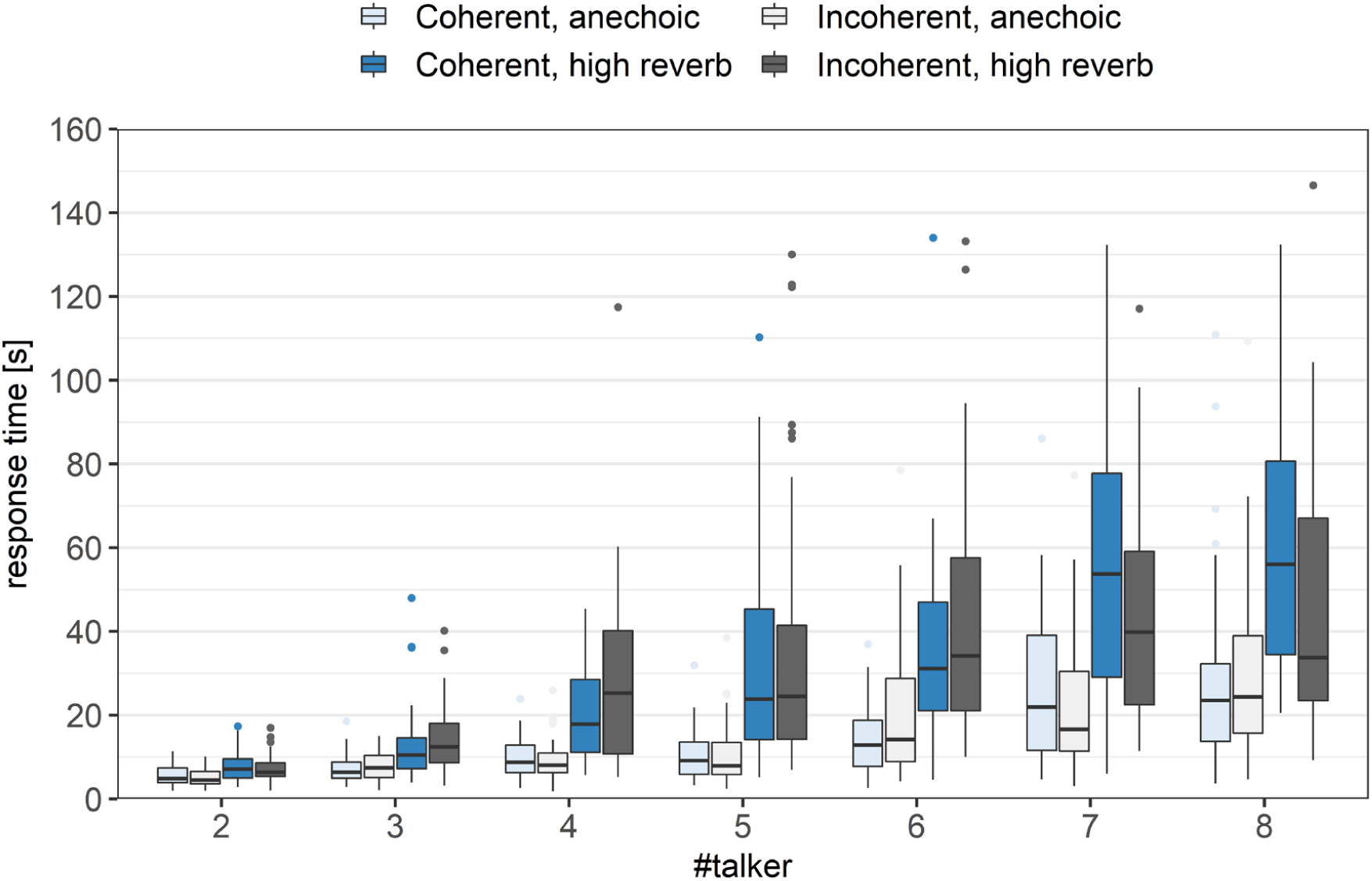
Response time with respect to the number of talkers in a scene. The light-blue and light-grey boxes indicate the anechoic room acoustic condition with coherent and incoherent visual information, respectively. The dark-blue and dark-grey boxes indicate the high reverberant room acoustic condition. The boxes cover the range between the 25th and the 75th percentile. The horizontal line in the boxes indicates the median. The whiskers extend to 1.5 times the inter-quartile range. Outliers are indicated as dots.

## IV. Discussion

In the current study we investigated the ability of normal-hearing listeners to identify and locate a story in the presence of other stories. The task of the listeners was to locate a target story in the presence of a varying number of simultaneous interfering talkers. Furthermore, the effect of audio-visual room information was investigated, by testing different audio-visually coherent and incoherent reverberant environments. The data showed that the localization accuracy and the response time are affected by the number of simultaneous talkers as well as by reverberation. With an increase of number of interfering talkers and an increase of reverberation time the performance of the listeners decreased. Presenting incoherent audio-visual room information did not affect the outcome measures.

### A. Effect of number of talkers

Several factors are likely to affect the increase in response time with increasing number of talkers. In the present study the speech level of each talker was kept constant independent of the number of talkers, and therefore, the signal-to-noise ratio (SNR) decreases. Thus, the intelligibility is expected to drop with the number of simultaneous talkers. However, the effective SNR is constantly changing with head-motion and fluctuations in the signals (Grange & Culling, 2016). The head-motion introduces a variation of the target and interferer angles relative to the head and thus head-shadow and interaural time differences vary. Both head-shadow and interaural time differences have been shown to be utilized to separate target and interfering speech sources (Bronkhorst, 2000; Culling et al., 2004). Fluctuations in the speech signals allow for dip-listening which can significantly improve the SNR in some time-frequency bins. Such glimpses can help to better understand speech (Glyde et al., 2013; Miller & Licklider, 1950). When many speech sources are presented, such glimpses are usually reduced (Cooke, 2006; Freyman et al., 2004).

Another effect that likely influences the response time is the amount of informational masking, i.e., confusions between the target and the interferers (Carhart et al., 1969; Durlach et al., 2003; Kidd et al., 2008; Watson, 2005). Previous studies have argued that the amount of informational masking decreases with increasing number of simultaneous talkers (Carhart et al., 1975; Freyman et al., 2004; S. A. Simpson & Cooke, 2005). However, in the current study the target speaker needs to be identified by understanding the speech and to do so, listeners also need to understand the content of the interferers. Thus, the listener needs to employ a strategy to search through the auditory scene and while performing the search an interfering talker becomes a temporary target talker. Therefore, the definition of informational masking that was already controversial in classic speech perception tasks (Durlach et al., 2003; Kidd et al., 2008; Watson, 2005) becomes even more complex. How the listeners perform this task and which search strategies they employ, remains an open question and is out of the scope of the current study.

### B. Effect of Reverberation

Reverberation was found to affect the response time only between the mid-reverberation and the high-reverberation conditions, and when there were four or more talker in a scene. In literature, it is reported that reverberation affects speech intelligibility more with few interfering talkers because potential speech gaps and pauses get ‘filled’ with the reverberant energy (Bolt & MacDonald, 1949; Xia et al., 2018). Such gaps generally do not exist with many overlapping speech sources (Cooke, 2006; Freyman et al., 2004). A potential explanation for the disagreement is that the task remains fairly easy with additional reverberation when few talkers are in a scene and thus, the effect of reverberation is masked.

No difference in response time was observed between the anechoic and the mid-reverberant conditions. The inexistent difference between the anechoic and the mid-reverberant condition contradicts results from previous studies where differences in speech perception between mildly reverberant conditions and anechoic conditions were found (Ahrens, Marschall, et al., 2019; Duquesnoy & Plomp, 1980; Plomp, 1976). The reason for this discrepancy could be that the test paradigm might not be as sensitive to capture small differences in reverberation time, as traditional speech tests. However, (Kopčo et al., 2010) discussed a similar finding that mild reverberation does not affect the speech localization in background speech by comparing their study with data from (B. D. Simpson et al., 2006). This raises the question if there is an effect of mild reverberation on speech intelligibility in everyday situations or if this effect can only be observed in artificial listening scenarios in the laboratory.

### C. Experimental paradigm

The spatial scene analysis method employed in this study was similar to (Weller et al., 2016). The most significant difference between the approaches is that in the current study the target speech stimulus needed to be understood while the task in (Weller et al., 2016) was to judge the gender of all talkers presented in a scene. Consequently, they used the total number of perceived talkers as their main outcome measure, while we used the response time. Furthermore, in their study the participants needed to translate the spatial percept from an egocentric auditory perception onto a top-down view interface. This translation was not needed in the current study as virtual reality was employed as a user interface.

While the use of virtual reality can allow for a more user-friendly interface, virtual reality could also introduce issues to an experiment. For example, the auditory percept might be affected by the physical presence of the headset which has been shown to be negligible for setups with far spaced sources (Ahrens, Lund, et al., 2019; Gupta et al., 2018). Furthermore, virtual reality glasses might alter the participant’s behavior due to their physical appearance but also because the visual world is not an exact copy of the real world. However, the influence is likely negligible in this experimental setup.

Contrary to classical speech perception studies where a %-correct or a reception threshold is determined, in the present study the response time was used as the main outcome measure. (Drullman & Bronkhorst, 2000) used a similar speech localization/identification paradigm with sentences and words instead of ongoing speech. They showed that the trend of change in intelligibility with increasing number of talkers was similar to the trend of the response times, i.e., with more interfering talkers the intelligibility decreases, and the response time increases. While the material and the task were not fully comparable between these studies, one can expect a correlation between speech intelligibility and response time.

### D. Effect of incoherent AV

Visual information is known to affect speech perception (McGurk & MacDonald, 1976). However, the effect of visual room information on auditory perception remains unclear. Previous studies showed that visual information of the room can improve auditory distance perception (Calcagno et al., 2012) and incongruent audio-visual cues can disrupt distance or externalization percepts (Gil-Carvajal et al., 2016). However, visual information has been shown to not affect the percept of reverberation (Schutte et al., 2019), which is in line with the results from the current study.

### E. Limitations

The speech material (10 stories spoken by 10 talkers) was recorded specifically for this study with the aim to have distinctly different content that can be visualized with an icon. Furthermore, we aimed for natural speech as opposed to highly controlled recordings with professional speakers. This approach also comes with disadvantage; for example some stories or talkers might be easier to understand than others. However, as stories and talkers were chosen randomly, their influence is likely to be little over the sufficiently large number of iterations.

One aim of this study was to develop a test paradigm that is more like real-life listening than most current speech intelligibility tests. While the task of understanding and locating a speech stream out of interfering speech is more similar to traditional speech tests, it is by no means a replications of a realistic cocktail-party situation. Firstly, all talkers are located at the same distance and with the same speech level and face the listener. This decision was made to not give any level, directional or direct-to-reverberant energy cues other than the information from the room reflections and the talkers themselves. Secondly, the visual avatars are highly conceptualized human bodies. Technology does not yet allow to visualize highly realistic human avatars with conventional computational power and effort. When using avatars that share similarities with real humans but evidently are not, viewers might get distracted (compare uncanny valley, (Diel et al., 2022)). Thirdly, lip-movements have not been included in this study. This choice was made because lip-movement simulations are not, as to the knowledge of the authors, evaluated for hearing research purposes. Additionally, the aim of the avatars was more to be a ‘response-box’ than an actual simulation of a human talker.

## V. Conclusions

In the present study we investigated the ability of listeners to analyze a spatial scene with multiple talkers. A varying number of simultaneously spoken stories was presented in different reverberant environments and listeners were asked to locate a target story. Results showed that the number of simultaneous talkers affected the correct identification as well as the response time. Reverberation only affected the outcome measures when the reverberation time was high but not with moderate reverberation.

## Acknowledgement

We would like to thank Marton Marschall, Valentina Zapata Rodriguez and Jakob Nygard Wincentz for their valuable feedback regarding the plausibility of the virtual audio-visual rooms and Torsten Dau for the feedback on the experimental design.

## Footnotes

1 Data available: https://data.dtu.dk/articles/Recordings_of_Danish_Monologues_for_Hearing_Research/9746285

## Notes

### Competing Interest Statement

The authors have declared no competing interest.

